# Genomic and anatomical comparisons of skin support independent adaptation to life in water by cetaceans and hippos

**DOI:** 10.1101/2020.11.15.383638

**Authors:** Mark S. Springer, Christian F. Guerrero-Juarez, Matthias Huelsmann, Matthew A. Collin, Kerri Danil, Michael R. McGowen, Ji Won Oh, Raul Ramos, Michael Hiller, Maksim V. Plikus, John Gatesy

## Abstract

The macroevolutionary transition from terra firma to obligatory inhabitance of the marine hydrosphere has occurred twice in the history of Mammalia: Cetacea and Sirenia. In the case of Cetacea (whales, dolphins, porpoises), molecular phylogenies provide unambiguous evidence that fully aquatic cetaceans and semiaquatic hippopotamids (hippos) are each other’s closest living relatives. Ancestral reconstructions further suggest that some adaptations to the aquatic realm evolved in the common ancestor of Cetancodonta (Cetacea+Hippopotamidae). An alternative hypothesis is that these adaptations evolved independently in cetaceans and hippos. Here, we focus on the integumentary system and evaluate these hypotheses by integrating new histological data for cetaceans and hippos, the first genome-scale data for pygmy hippopotamus, and comprehensive genomic screens and molecular evolutionary analyses for protein-coding genes that have been inactivated in hippos and cetaceans. We identified ten skin-related genes that are inactivated in both cetaceans and hippos, including genes that are related to sebaceous glands, hair follicles, and epidermal differentiation. However, none of these genes exhibit inactivating mutations that are shared by cetaceans and hippos. Mean dates for the inactivation of skin genes in these two clades serve as proxies for phenotypic changes and suggest that hair reduction/loss, the loss of sebaceous glands, and changes to the keratinization program occurred ~16 million years earlier in cetaceans (~46.5 Ma) than in hippos (~30.5 Ma). These results, together with histological differences in the integument and prior analyses of oxygen isotopes from stem hippopotamids (“anthracotheres”), support the hypothesis that aquatic adaptations evolved independently in hippos and cetaceans.

## INTRODUCTION

The long evolutionary history of Mammalia is mostly one of deployment and adaptation to different terrestrial habitats. Nevertheless, numerous mammalian clades have returned to aquatic habitats, either on a part-time basis or with full commitment to the aquatic realm. Fully aquatic clades include Cetacea and Sirenia, and in both cases there are extinct taxa that document the macroevolutionary transition from land to water [1,2]. Semiaquatic clades are much more numerous and examples are found in a wide range of mammalian orders such as Monotremata (platypuses), Didelphimorphia (water opossums), Afrosoricida (otter shrews), Rodentia (muskrats, beavers, capybaras), Eulipotyphla (desmans, star-nosed moles), Carnivora (pinnipeds, otters, mink), and Cetartiodactyla (hippos). Adaptations to aquatic habitats are most extreme in fully aquatic forms where virtually every organ system has been highly modified in conjunction with the transition to life in the water. For example, locomotory adaptations in cetaceans include hind limb loss, modification of the front limbs into flippers, and conversion of the tail into a powerful fluke [1]. At the level of sensory systems, cetaceans have highly modified eyes with reduction or loss of color vision [3], olfactory structures that are highly reduced or absent [4,5], and in the case of odontocetes (toothed whales) ultrasonic hearing that enables echolocation [6].

Cetaceans also show major changes to their integument. By contrast with terrestrial mammals, the skin of fully aquatic mammals constantly interacts with dense, viscous and highly thermally conductive water, which poses unique physical challenges to its outer surface. The epidermis of cetaceans is unusually thick and undergoes constant cellular renewal [7–13]. Even with its increased thickness there are only three distinct histological layers – a basal layer (*stratum basale*), an intermediate layer (*stratum spinosum*), and an outer layer (*stratum corneum*) [14]. The *stratum granulosum*, which in land mammals lies beneath the *stratum corneum,* is either ill-defined or absent [10,15]. The exceptionally tall epidermis provides additional mechanical and thermal protection, and its fast rate of sloughing and extensive epidermal-dermal interdigitations guard against potential damage from shear stress during locomotion [16]. The *stratum basale* forms deep root-like projections, known as rete ridges, that extend into the underlying dermis [17–19]. Extensive cytoplasmic lipid vacuoles are also present in the *stratum spinosum* and *stratum corneum* keratinocytes [20–22], and have been speculated to play metabolic and/or thermo-insulating roles [22,23]. Unlike in terrestrial mammals, cells of the *stratum corneum* retain nuclei and do not become fully keratinized in cetaceans [14]. This is likely due to fully aquatic mammals not needing a functional epidermal barrier to a dry environment [24]. The cetacean dermis is also thickened and consists of an upper papillary layer that intricately interdigitates with the epidermis and a lower reticular layer that gradually transitions into an adipose layer called blubber [10,11,18,25,26].

Cetaceans have dramatically reduced the diversity, density, and distribution of ectodermal appendages including hair follicles and skin glands [27]. They show no evidence of pelage (i.e., fur type) hair follicle formation during embryonic development although vibrissa follicles (i.e., whiskers) form on the head [28–32]. Mysticetes and some odontocetes cyclically grow whisker hairs as adults, but most odontocetes lose their whiskers as adults and convert vibrissa follicles into degenerated pits (i.e., crypts) thought to perform sensory functions [30,33,34]. Typical hair follicles in terrestrial mammals are associated with oil-secreting sebaceous glands, but these are lacking in the published histology of cetacean vibrissa follicles [31,32,35]. Cetaceans also lack sweat glands [1,14,27].

Among extant semiaquatic forms, hippos are the largest herbivores [36]. Both extant species (*Hippopotamus amphibius* [river hippo], *Choeropsis liberiensis* [pygmy hippo]) are nocturnal. Hippos spend their days in the water where they remain cool, but emerge at dusk to feed on grasses and other vegetation. *H. amphibius* and *C. liberiensis* have superficially similar-looking skin, with histological studies primarily on the former [15,37,38] with one recent exception [39]. Similar to cetaceans, adult *H. amphibius* has a thick epidermis and its *stratum basale* forms distinct projections into the papillary dermis [15]. *H. amphibius* also shows elevated levels of lipid storage in the epidermis. However, as compared to cetaceans, most prominent lipid deposits occur in intercellular locations within the *stratum corneum* [23]. Hippos have numerous bristle-like whiskers on their muzzle and tail, but unlike cetaceans they also have pelage hairs that are sparsely distributed across most of the body [15]. Sebaceous glands have not been reported in hippos, but previous histological studies have only examined limited regions of the body [15,39]. Hippo skin contains anatomically complex sweat glands [37,38] that secrete a distinct red-orange pigmented sweat that is thought to have sunscreen and/or antimicrobial properties [40,41].

The traditional view based on phylogenetic interpretations of morphology is that cetaceans are excluded from a monophyletic Artiodactyla (even-toed hoofed mammals) [42,43] and that hippopotamids are the sister taxon to pigs (Suidae) and/or peccaries (Tayassuidae) [44–48]. However, molecular studies challenged this view and eventually provided conclusive evidence that cetaceans are nested within Artiodactyla [49,50], specifically as the sister taxon to Hippopotamidae [51–54].

Given that cetaceans and hippopotamids share a variety of morphological and behavioral characters that may be related to aquatic habitats (hairless or nearly hairless body, lack of sebaceous glands, lack of scrotal testes, underwater parturition and nursing of young, underwater detection of sound directionality), the most parsimonious hypothesis is that these shared characters evolved in the common ancestor of these two clades (= Cetancodonta) and that this common ancestor was semiaquatic [51,52,55,56] (Figure 1A). O’Leary and Gatesy [57] argued in favor of this common origin hypothesis based on ancestral reconstructions of underwater hearing and other aquatic characters. Gatesy et al. [1] favored the aquatic ancestry hypothesis, specifically in freshwater, based on oxygen isotope values in pakicetid cetaceans, the presence of dense, osteosclerotic bones in both hippos and basal stem cetaceans (raoellids, pakicetids), as well as shared behavioral and soft-anatomical traits. Alternatively, semiaquatic adaptations may have evolved convergently in hippos and cetaceans [36] (Figure 1B). In part, support for these two competing hypotheses turns on the phylogenetic placement of various ‘Anthracotheriidae’, which collectively are the paraphyletic stem group to Hippopotamidae. Some anthracotheres in the subfamilies Anthrocotheriinae (e.g., *Anthracotherium*) and Microbunodontinae (e.g., *Microbunodon*) are inferred to have been terrestrial based on oxygen isotope values, but members of Bothriodontinae (e.g., *Bothriogenys*, *Merycopotamus*, *Libycosaurus*) have values that are consistent with a semiaquatic lifestyle [58–62]. Cooper et al. [62] performed ancestral reconstructions on the microanatomy of the transitional zone between the medullary cavity and the cortical compacta of the tibia for a data set that included both anthracotheres and stem cetaceans (“archaeocetes”) and concluded that the most recent common ancestor of Cetancodonta was probably semiaquatic. More recently, Soe et al. [63] reported new skeletal material for the Eocene anthracotheriid *Siamotherium*, which they recovered as the most basal genus of anthracotheres in their phylogenetic analysis. Soe et al. [63] concluded that there is no evidence of clear-cut aquatic adaptations in *Siamotherium* fossils. These authors did not perform ancestral-state reconstructions, but the successive placement of two presumably terrestrial anthracothere clades (*Siamotherium*, Anthracotheriinae) at the base of Hippopotamoidea suggests that aquatic specializations in hippopotamids and cetaceans are convergent.

**Figure 1.**
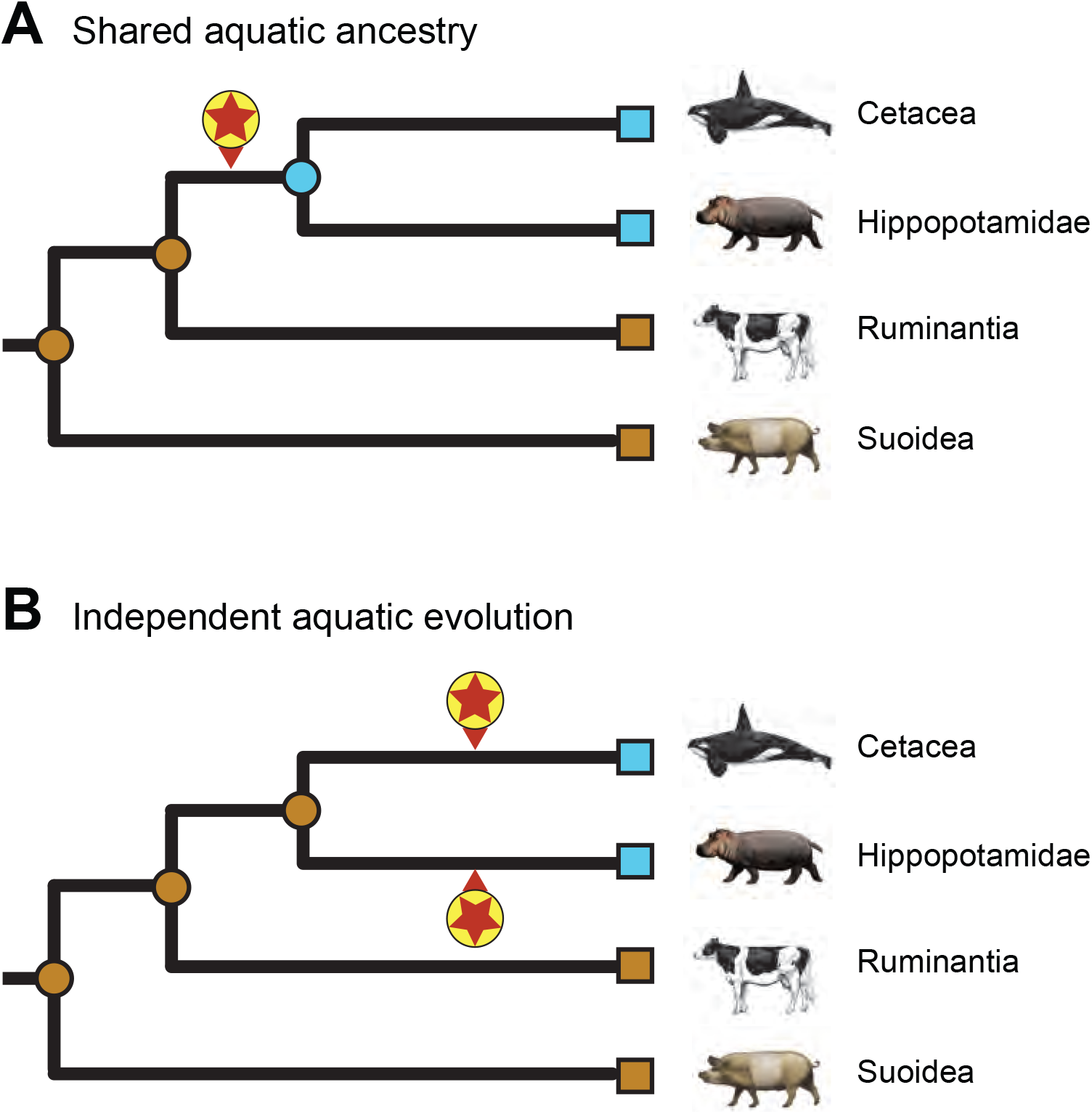
Two hypotheses for the evolution of aquatic adaptations in hippopotamids and cetaceans. (A) Evolution of shared aquatic features in the common ancestor of Hippopotamidae and Cetacea, and (B) independent evolution of aquatic features on the cetacean stem lineage and also on the hippopotamid stem lineage. Encircled red stars mark the initial evolution of behavioral, physiological, and anatomical characteristics associated with adaptation to aquatic environments. Aquatic (blue) and terrestrial (brown) specializations of extant taxa (squares) and ancestral nodes (circles) are indicated in each scenario.

At the molecular level, positive selection analyses on genomic/transcriptomic data provide minimal support for shared aquatic adaptations in the ancestry of Cetancodonta [64]. However, given that most aquatic similarities between cetaceans and hippos are in the skin and that branch-site selection analyses can have weak statistical power, a more promising approach may be to search for skin genes that have been inactivated in the common ancestor of cetaceans and hippos that relate to aquatic specialization of the skin. Previous authors have successfully employed this approach to identify numerous skin genes that were inactivated in the common ancestor of cetaceans, either with a candidate-gene approach [16,24,65] or with comprehensive genomic screens [66–68]. To date, only two candidate gene studies [69–70] have searched for candidate skin genes that are inactivated in cetaceans and hippos. Springer and Gatesy [69] reported that two skin-related genes, *MC5R* and *PSORS1C2*, are inactivated in cetaceans but not hippos. Lopes-Marques et al. [70], in turn, reported independent inactivation of two different sebaceous gland genes (*AWAT1*, *MOGAT3*) in hippos and cetaceans. Hence, it remains to be determined if a comprehensive screen [*sensu* 67] will reveal additional skin genes that have been inactivated in both cetaceans and hippos.

Here, we present new morphological and molecular evidence to evaluate competing hypotheses that aquatic adaptations evolved in the common ancestor of cetaceans and hippos versus independently in these two groups (Figure 1). First, we provide histological data for different regions of the integument in two cetacean species (*Tursiops truncatus* [bottlenose dolphin], *Eschrichtius robustus* [gray whale]) and both extant hippos (*Hippopotamus amphibius* [river hippo], *Choeropsis liberiensis* [pygmy hippo]). Next, we perform a comprehensive screen of genomes from representative cetaceans and hippopotamids (including new Illumina data for the pygmy hippopotamus) for protein-coding genes that have been inactivated in both of these clades. We then assess the timing of pseudogenization for these genes based on the presence/absence of shared inactivating mutations and selection intensity (dN/dS) analyses that are integrated with a molecular timetree. Finally, we synthesize our molecular and histological findings with previous studies and discuss these results in the context of the rich fossil record for stem cetaceans (“archaeocetes”) and stem hippos (“anthracotheres”).

## RESULTS AND DISCUSSION

### Comparative Skin Histology Between Terrestrial and Aquatic Cetartiodactyla

We analyzed skin samples from numerous body regions (face, eyelid, ear, dorsum, ventrum and tail) in both hippopotamid species, and from the snout region of one odontocete (bottlenose dolphin) and one mysticete (gray whale). Table 1 summarizes features of the skin in these taxa and also includes data from the literature for terrestrial mammals including humans and two cetartiodactyls (cow, pig) that are close relatives of Cetancodonta [71–79].

**Table 1.**
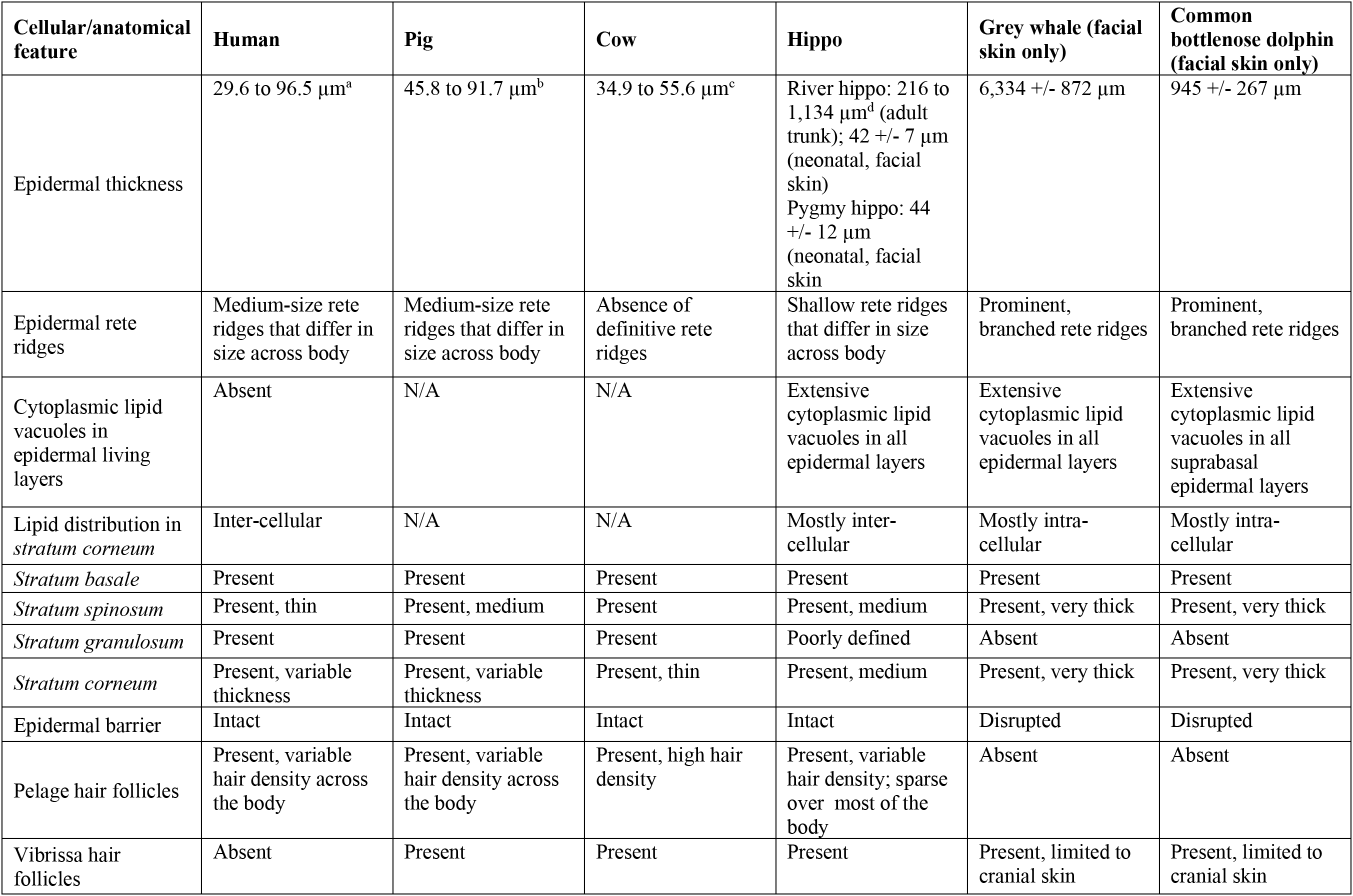
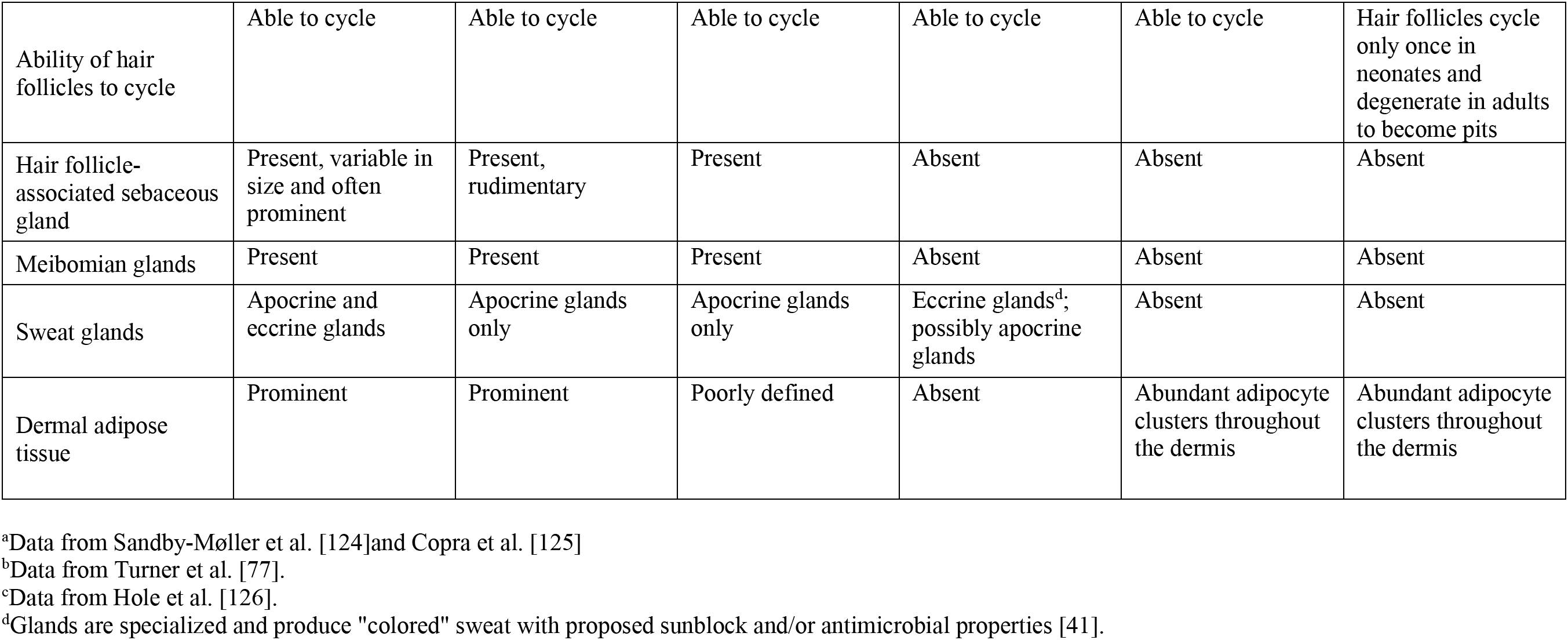
Features of the skin in human, cow, pig, hippo, and two cetaceans.

| Cellular/anatomical feature | Human | Pig | Cow | Hippo | Grey whale (facial skin only) | Common bottlenose dolphin (facial skin only) |
| --- | --- | --- | --- | --- | --- | --- |
| Epidermal thickness | 29.6 to 96.5 $\mu\text{m}^{\text{a}}$ | 45.8 to 91.7 $\mu\text{m}^{\text{b}}$ | 34.9 to 55.6 $\mu\text{m}^{\text{c}}$ | River hippo: 216 to 1,134 $\mu\text{m}^{\text{d}}$ (adult trunk); 42 +/- 7 $\mu\text{m}$ (neonatal, facial skin)<br>Pygmy hippo: 44 +/- 12 $\mu\text{m}$ (neonatal, facial skin) | 6,334 +/- 872 $\mu\text{m}$ | 945 +/- 267 $\mu\text{m}$ |
| Epidermal rete ridges | Medium-size rete ridges that differ in size across body | Medium-size rete ridges that differ in size across body | Absence of definitive rete ridges | Shallow rete ridges that differ in size across body | Prominent, branched rete ridges | Prominent, branched rete ridges |
| Cytoplasmic lipid vacuoles in epidermal living layers | Absent | N/A | N/A | Extensive cytoplasmic lipid vacuoles in all epidermal layers | Extensive cytoplasmic lipid vacuoles in all epidermal layers | Extensive cytoplasmic lipid vacuoles in all suprabasal epidermal layers |
| Lipid distribution in *stratum corneum* | Inter-cellular | N/A | N/A | Mostly inter-cellular | Mostly intra-cellular | Mostly intra-cellular |
| *Stratum basale* | Present | Present | Present | Present | Present | Present |
| *Stratum spinosum* | Present, thin | Present, medium | Present | Present, medium | Present, very thick | Present, very thick |
| *Stratum granulosum* | Present | Present | Present | Poorly defined | Absent | Absent |
| *Stratum corneum* | Present, variable thickness | Present, variable thickness | Present, thin | Present, medium | Present, very thick | Present, very thick |
| Epidermal barrier | Intact | Intact | Intact | Intact | Disrupted | Disrupted |
| Pelage hair follicles | Present, variable hair density across the body | Present, variable hair density across the body | Present, high hair density | Present, variable hair density; sparse over most of the body | Absent | Absent |
| Vibrissa hair follicles | Absent | Present | Present | Present | Present, limited to cranial skin | Present, limited to cranial skin |
| Ability of hair follicles to cycle | Able to cycle | Able to cycle | Able to cycle | Able to cycle | Able to cycle | Hair follicles cycle only once in neonates and degenerate in adults to become pits |
| Hair follicle-associated sebaceous gland | Present, variable in size and often prominent | Present, rudimentary | Present | Absent | Absent | Absent |
| Meibomian glands | Present | Present | Present | Absent | Absent | Absent |
| Sweat glands | Apocrine and eccrine glands | Apocrine glands only | Apocrine glands only | Eccrine glands<sup>d</sup>; possibly apocrine glands | Absent | Absent |
| Dermal adipose tissue | Prominent | Prominent | Poorly defined | Absent | Abundant adipocyte clusters throughout the dermis | Abundant adipocyte clusters throughout the dermis |

<sup>a</sup>Data from Sandby-Møller et al. [124]and Copra et al. [125]
<sup>b</sup>Data from Turner et al. [77].
<sup>c</sup>Data from Hole et al. [126].
<sup>d</sup>Glands are specialized and produce "colored" sweat with proposed sunblock and/or antimicrobial properties [41].

Hippos and cetaceans have prominent differences in the thickness and organization of the epidermis (Figure 2). Consistent with previous reports [17–19], the facial epidermis in adult *Eschrichtius robustus* (Figure 2E) and neonatal *Tursiops truncatus* (Figure 2F-G) is very thick, with a wide *stratum spinosum*, and an undulated *stratum basale* with deep root-like rete ridges that extend into the dermis. By contrast, the epidermis in neonatal pygmy hippo is thin, with only shallow rete ridges (Figure 2A-D). Among different anatomical sites, epidermal rete ridges are most prominent in the lip and tail and least prominent in the ear and ventral skin.

**Figure 2.**
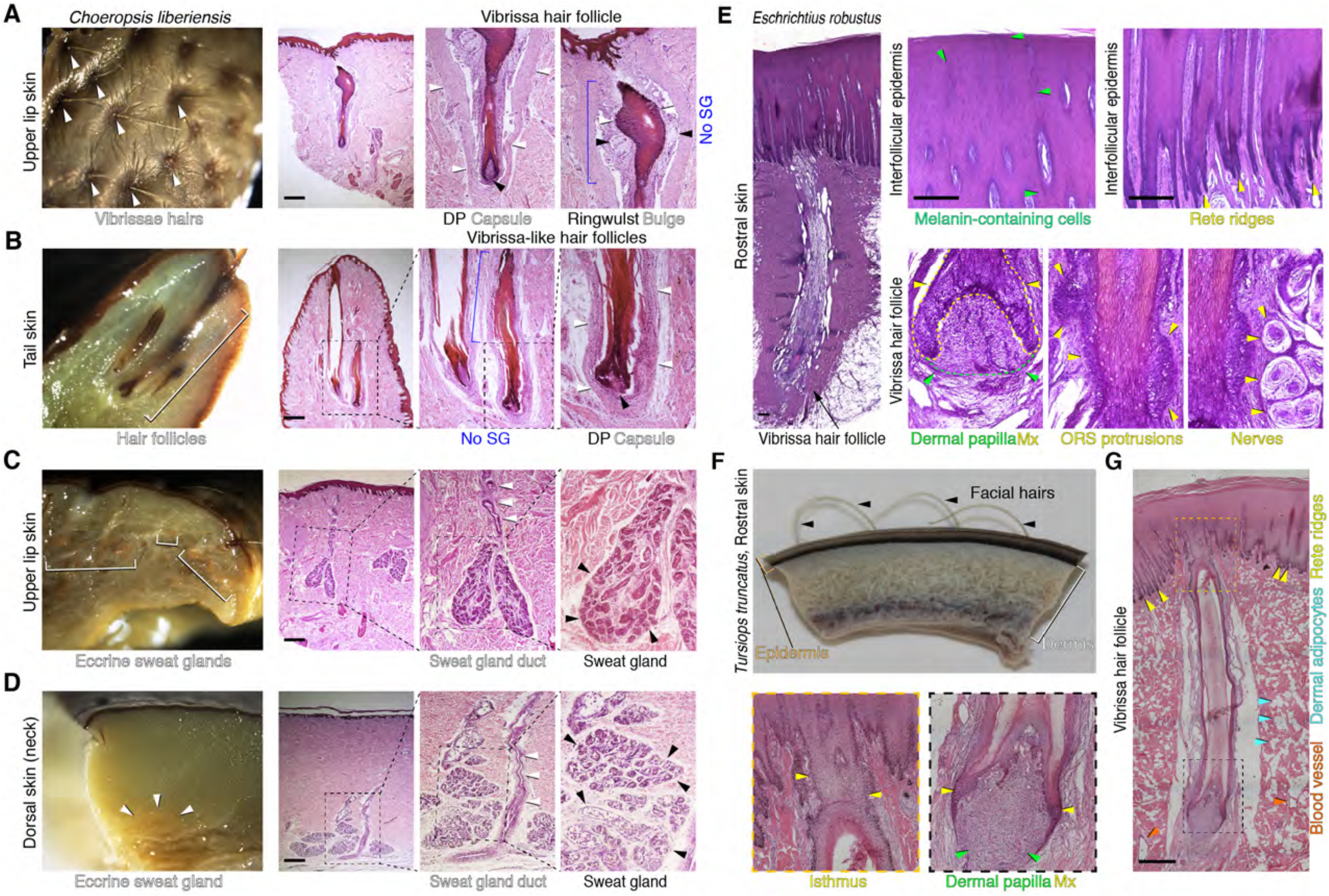
Histological features of the skin in hippos (A-D) and cetaceans (E-G). (A) Whole mount (left) and histological appearance (middle, right) of the upper lip skin in pigmy hippo. In the lip, vibrissa hairs above the skin surface and anagen (active growth) phase vibrissa hair follicles are prominent. On histology, each lip vibrissa follicle has a prominent mesenchymal dermal papilla (black arrowhead, center), a collagen capsule (white arrowheads, center), an epithelial matrix, a mesenchymal ringwulst (black arrowheads, right), and an epithelial bulge (white arrowheads, right). There is no histological evidence of sebaceous glands. (B) Whole mount of tail skin from pygmy hippo (left) shows large hair follicles (white bracket). Histological analysis (right) suggests that tail hair follicles might be of vibrissa type because they are surrounded by a collagen capsule (white arrowheads). There is no histological evidence of sebaceous glands. (C, D) Whole mount and corresponding histological view of upper lip skin (C) and dorsal skin (D) in the pigmy hippo. At both sites, eccrine sweat glands are present. On histology, secretory coils located deep in the dermis are marked with black arrowheads; associated excretory ducts (where obvious) are marked with white arrowheads. In the dorsal skin (D), secretory coils of the glands reside at the very base of the dermis and come in contact with the underlying skeletal muscle layer. There is no histological evidence of dermal adipocytes. (E) Histology of facial skin and a rostral vibrissa hair follicle in an adult gray whale. The epidermis is thick and its basal layer is heavily undulated. The vibrissa follicle has a typical anagen (active growth) phase morphology with a large mesenchymal dermal papilla (green arrowheads, second panel) surrounded by epithelial matrix (yellow arrowheads, second panel). The epithelial outer root sheath compartment located above hair matrix is uncharacteristically thick and has prominent protrusions (yellow arrowheads, third panel). The vibrissa follicle is associated with distinct nerve bundles on either side (yellow arrowheads, fourth panel). (F) Wholemount side view of rostral skin from neonatal common bottlenose dolphin. Vibrissae hairs (black arrowheads) are clearly visible above the skin surface. (G) Histological view of rostral vibrissa hair follicle from neonatal common bottlenose dolphin. The vibrissa follicle has anagen phase morphology. A large dermal papilla (green arrowheads, bottom middle panel) and an uncharacteristically thin epithelial matrix (yellow arrowheads, bottom middle panel) are obvious. Unlike in gray whale, the outer root sheath lacks undulations, and like in gray whale, the vibrissa follicle lacks sebaceous gland. The overlaying epidermis displays prominent rete ridges (yellow arrowheads, right panel). The surrounding dermis contains numerous adipocyte clusters (blue arrowheads, right panel). Scale bar: A-E, G–50 μm.

Hippo skin contains hair follicles of both pelage and vibrissa morphology. Prominent vibrissa follicles that contain collagen capsules and ringwulst (ring-like dermal structure that surrounds the follicle) were observed in the upper lip skin (Figure 2A; Supplemental Figure S1C). Some tail-tip hair follicles also display collagen capsules (Figure 2B; Supplemental Figure S1E), suggesting that the tail region might contain a mixture of pelage and vibrissa hair types. Hair follicles in ear and eyelid skin have typical pelage morphology, and meibomian glands are absent in the latter (Figure 2D; Supplemental Figure S1A-B, S2C). In all samples studied, hippo hair follicles lack hair-associated sebaceous glands (Figure 2; Supplemental Figure S1). No hair follicles were identified in the available dorsal and ventral skin samples. The facial skin of adult gray whale and neonatal bottlenose dolphin had prominent vibrissa hair follicles (Figure 2E-G), yet their structure differs from that of facial hippo vibrissae. Cetacean vibrissae lack both collagen capsules and ringwulst. In the baleen whale (Figure 2E), but not the dolphin (Figure 2F-G), the epithelial outer root sheath is highly undulated and peri-follicular nerves are very distinct. In both species, mesenchymal dermal papillae and the epithelial hair matrix, two defining structures of actively growing hair follicles, are present. However, the hair matrix in dolphin vibrissae is uncharacteristically thin. There is no evidence of vibrissa-associated sebaceous glands in either cetacean species that was examined.

Prominent sweat glands are present in pygmy hippo skin in several locations, including the upper lip and both dorsal and ventral skin (Figure 2C-D; Supplemental Figure S2). The structure of the dermis is different in cetaceans and hippos. There are abundant clusters of adipocytes throughout the dermis in *Tursiops truncatus* (Figure 2G), but we found no histological evidence for mature adipocytes in hippo skin at all locations studied. In hippos, there are prominent differences in dermal thickness across body sites, with tail and ear dermis being the thinnest.

In conclusion, hippo skin is characterized by an epidermis that is thinner than in cetaceans, shallow rete ridges, a dermis of variable thickness without adipocytes, highly specialized sweat glands, and both pelage and vibrissa hair follicles. Sweat glands and both types of hair follicles display differing regional distributions. In contrast, cetacean skin has a distinctly thick and highly undulated epidermis, a thick and adipocyte-rich dermis, and few morphologically-specialized facial vibrissa hair follicles as the only skin appendage. Unlike humans and terrestrial cetartiodactyls, hair-associated sebaceous glands are lacking in hippos and cetaceans (Table 1).

### Genomic Screens and Patterns of Gene Inactivation in Cetaceans and Hippos

Next, we investigated the hypothesis of shared versus independent aquatic ancestry using existing and newly-generated genomic data. We first screened genomic alignments with 63 mammalian taxa for protein-coding genes that are inactivated in Hippopotamidae and Cetacea but not in terrestrial cetartiodactyls. Our initial screen included *Hippopotamus amphibius*, one baleen whale (*Balaenoptera acutorostrata* [common minke whale]), and three toothed whales (*Physeter macrocephalus* [sperm whale]*, Orcinus orca* [killer whale]*, Tursiops truncatus* [bottlenose dolphin]). Based on these screens, we identified 38 genes that have inactivating mutations (frameshift insertions/deletions, premature stop codons, splice site mutations, deleted exons) or are completely deleted in these taxa (Supplemental Table S1) including ten genes that have primary or sole functions related to skin and its ectodermal appendages (*ALOX15*, *AWAT1*, *KPRP*, *KRT2*, *KRT26*, *KRT77*, *KRTAP6-2*, *KRTAP6-3*, *KRTAP7-1*, *TCHHL1*). Two genes (*KRTAP6-2*, *KRTAP6-3*) were removed from this list because of ambiguous orthology relationships, leaving eight genes for a detailed analysis. We added *ABCC11* to this list, which is a candidate gene of interest that is expressed in axillary sweat glands in humans [80,81].

Since base errors in genome assemblies can mimic gene-inactivating mutations [82], we first used additional genomic data to confirm the validity of inactivating mutations and thus the loss of function of the nine genes. To this end, we used BLAST to investigate whether inactivating mutations are shared with independently-sequenced and assembled sister species genomes, which would indicate that these mutations are real and occurred in the common ancestor of taxa sharing the mutation(s). Our investigation of ten additional cetacean genomes revealed that all nine genes have mutations shared between at least two species, as exemplified in Figure 3A-B. Three of these nine genes (*AWAT1*, *KRTAP7-1*, *ABCC11*) exhibit inactivating mutations shared between odontocetes and mysticetes, indicating gene loss in the stem cetacean lineage (Figure 3C). Two genes (*KRT26*, *KRT77*) exhibit large or entire gene deletions in odontocetes and mysticetes, and gene loss dating (below) indicates that both losses happened in the stem cetacean lineage. The distribution of inactivating mutations in *ALOX15* suggests that it was independently inactivated on the stem branch of Delphinida, on the stem branch of Physeteridae (sperm whales; two stop codons are shared between *Physeter macrocephalus* and *Kogia breviceps*), and in the common ancestor of *Balaenoptera acutorostrata* and *B. bonaerensis* (minke whales). Finally, three genes are absent from all cetacean (*KPRP* and *TCHHL1*) or all odontocete (*KRT2*) genomes,. However, large deletions and rearrangements in these loci obscure reconstructions of when these genes were lost because there are no obvious breakpoints that are shared by odontocetes and mysticetes.

**Figure 3.**
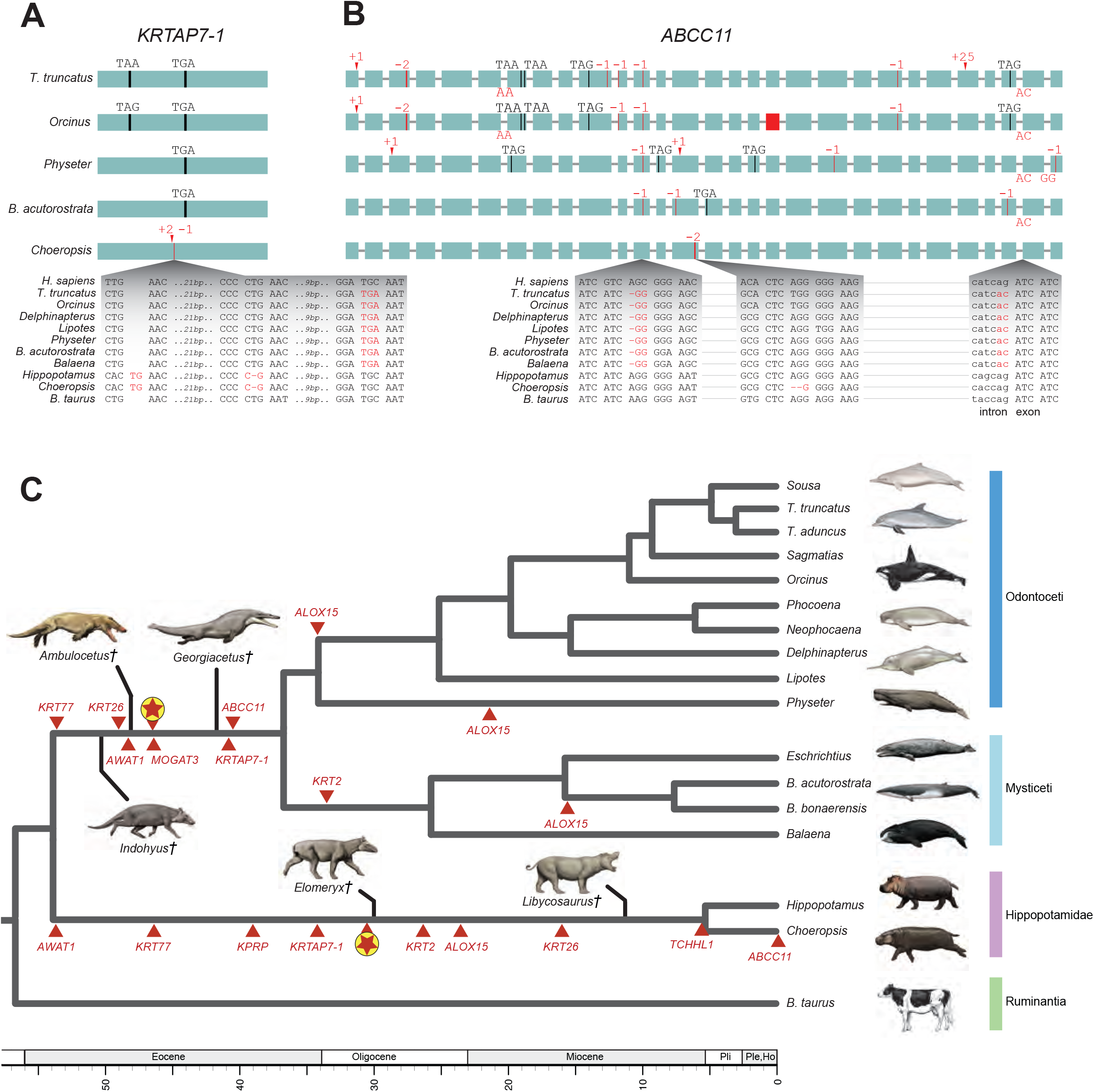
Inactivating mutations in *KRTAP7-1* (A) and *ABCC11* (B). Genes are shown with exons represented by green rectangles proportional to their size and introns represented by horizontal lines. Inactivating mutations are premature stop codons (vertical black line and corresponding triplet), insertions (red arrowhead and corresponding insertion size), deletions (vertical red line and corresponding deletion size, or red rectangle for completely deleted exon), and splice site donor or acceptor mutations (red letters at the end or beginning of an exon, respectively). Insets show the DNA sequence context of representative mutations. See supplementary alignment files for inactivating mutations in other genes. (C) The phylogenetic pattern of skin gene inactivations mapped onto a timetree for Cetancodonta (Hippopotamidae + Cetacea). Independent inactivations of skin genes (red triangles) are marked on branches of the tree; inactivation times are average estimates for each locus based on dN/dS analyses. The mean dates of skin gene knockouts for six loci that were inactivated in the common ancestor of Cetacea and for eight loci that were inactivated in the common ancestor of Hippopotamidae are indicated on the stem lineages to these clades (encircled red stars). The timetree for extant lineages is based on the molecular clock analysis of McGowen et al. [123]. Extinct lineages for stem cetaceans (*Indohyus*, *Ambulocetus*, *Georgiacetus*) and stem hippopotamids (*Elomeryx*, *Libycosaurus*) are approximately positioned relative to the geological time scale based on earliest occurrence in the fossil record for each genus and phylogenetic hypotheses based on morphological characters.

Genomic data for *Choeropsis liberiensis* are required to investigate whether mutations are shared between both extant hippopotamid species. Therefore, we generated ~40X of Illumina whole genome shotgun data of *C. liberiensis* and mapped these reads to the *Hippopotamus amphibius* assembly to obtain orthologous sequences. We found that both hippopotamids exhibit shared inactivating mutations in eight of these nine genes (*ALOX15*, *AWAT1*, *KPRP*, *KRT2*, *KRT26*, *KRT77*, *KRTAP7-1*, *TCHHL1*) (Figure 3). *ABCC11* is intact in *H. amphibius*, but we detected a two-bp frameshift deletion in exon 14 of *C. liberiensis*, suggesting a recent loss in pygmy hippo (Figure 3B).

Next we analyzed whether these nine skin-related gene losses have inactivating mutations that are shared by hippos and cetaceans, which would indicate gene inactivation in the common ancestral lineage of extant cetancodontans. We did not find any evidence of inactivating mutations that are shared by hippos and cetaceans. Instead, all nine genes have different mutations in cetaceans and hippopotamids (Figure 3, Supplemental Table S2), indicating that these genes were independently lost in both lineages. In a previous study on candidate sebaceous gland genes, Lopes-Marques et al. [70] reported the convergent loss of both *AWAT1* and *MOGAT3* in Cetacea and *Hippopotamus amphibius*. Our results confirm the independent inactivation of *AWAT1* in these two clades and further suggest that *AWAT1* was inactivated in the common ancestor of *H. amphibius* and *Choeropsis liberiensis*. However, our investigation of *MOGAT3* revealed that this gene underwent a tandem duplication, most likely in the common ancestor of the two extant hippopotamids (Figure 4). The pseudogenized copy of *MOGAT3* reported by Lopes-Marques et al. [70] is ~15.7 kb upstream of the functional copy of this gene on NCBI contig PVJP02910399. *C. liberiensis* retains an intact copy of *MOGAT3*, but we were unable to find evidence for retention of the pseudogenic paralog in this species. Lopes-Marques et al.’s [70] analyses of a *de novo* skin transcriptome provide additional support for two copies of *MOGAT3* in *H. amphibius*. One represents transcription from the intact *MOGAT3* locus whereas the other represents transcription of the inactivated copy of *MOGAT3*.

**Figure 4.**
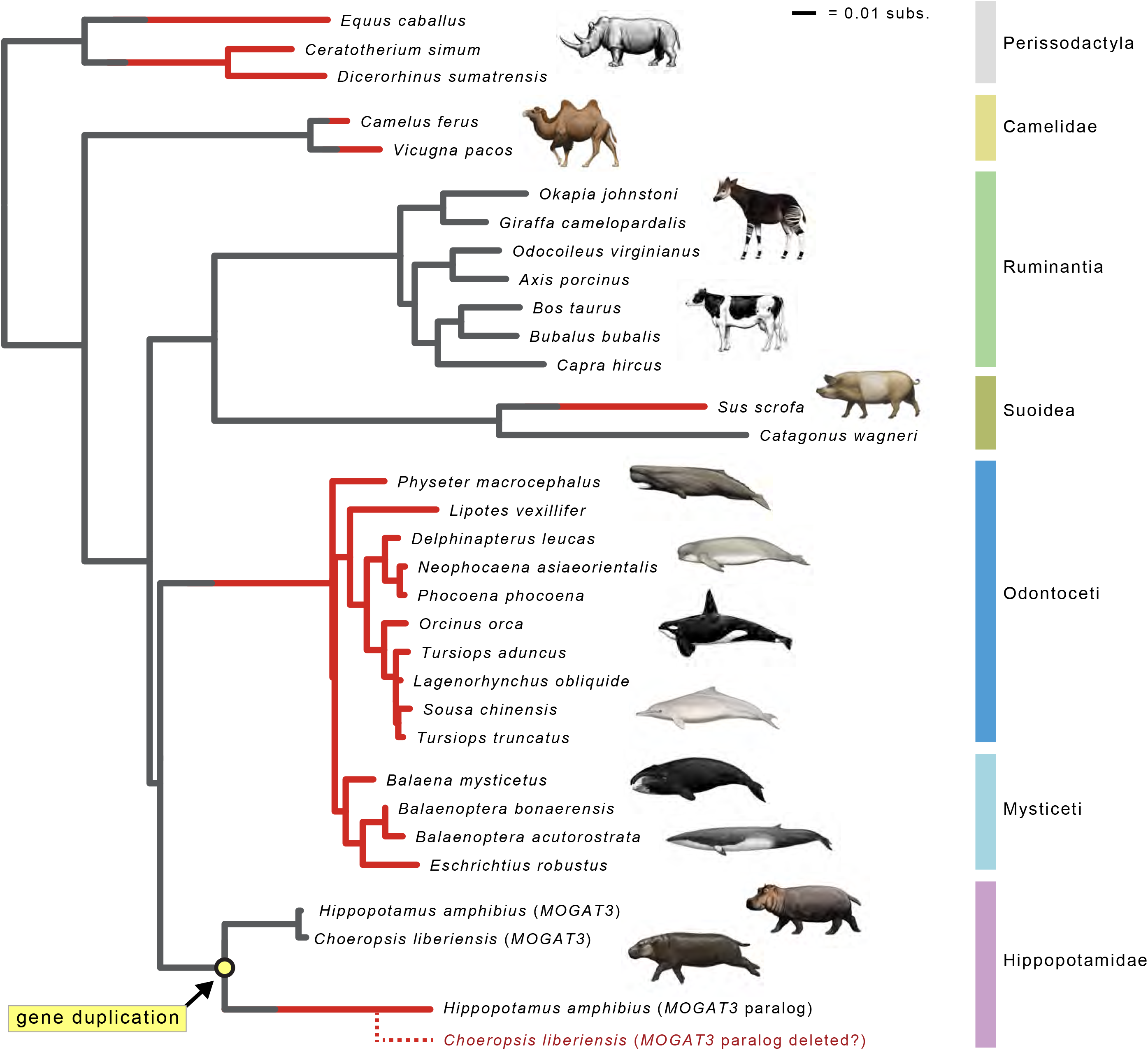
Gene tree for *MOGAT3* that shows inferred gene duplication event in Hippopotamidae (yellow circle) and parallel gene inactivations (red lineages). Seven gene knockouts are inferred, including pseudogenization of the *MOGAT3* paralog that was derived from a duplication event on the stem hippopotamid branch. Lineages with functional *MOGAT3* are gray, and lineages with inactivated *MOGAT3* are colored red. Branches where inactivation events (frameshift indels, premature stop codons) were inferred by parsimony optimization of indels are gray and red, indicating the transition from functional to non-functional. The dashed red lineage for *Choeropsis liberiensis* represents hypothesized deletion of the *MOGAT3* paralog in the genome of this species. Branch lengths are in expected numbers of substitutions per site.

### Comparison of Inactivated Skin Genes and Epidermal Phenotypes in Cetaceans and Hippos

The cetacean epidermis is very thick and renews rapidly, yet does not fully differentiate into the highly keratinized *stratum corneum* of terrestrial mammals that confers barrier functions. At the genetic level, these modifications are associated with loss of function of numerous genes that are part of the Epidermal Differentiation Complex (EDC), including S100 fused-type protein (*CRNN, FLG* [in mysticetes and sperm whales], *FLG2, HRNR, RPTN, TCHH, TCHHL1*, *TCHHL2*) [27,65] and suprabasal epidermal keratin genes (*KRT1, KRT2, KRT9, KRT10, KRT77, KRT23* [odontocetes only], *KRT24*) [16]. Our analyses on cetacean genomes confirm these findings (Figure 5).

**Figure 5.**
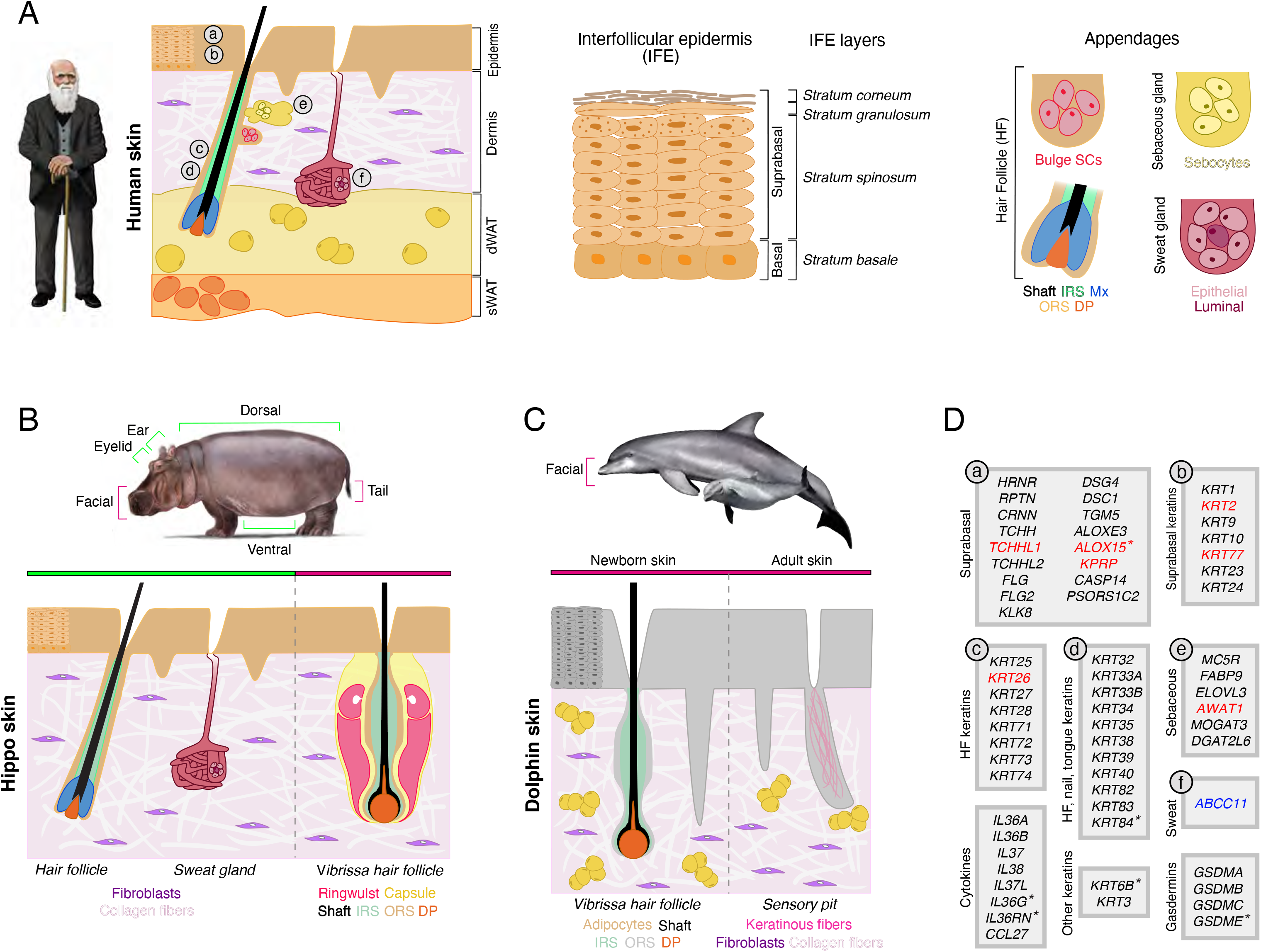
A comparison of skin structures in human, hippo, and dolphin with skin-associated gene inactivation in hippopotamids and cetaceans. A) Schematic drawing of *Homo sapiens* skin. Key anatomical structures, including multi-layered epidermis, dermis, dermal white adipose tissue (dWAT), subcutaneous white adipose tissue (sWAT), and ectodermal appendages (hair follicles with sebaceous glands and sweat glands) are shown and color-coded. The epidermis (middle panel) is divided into *stratum basale* – which houses stem cells, and suprabasal layers of differentiating cells that include *stratum spinosum*, *stratum granulosum*, and *stratum corneum.* dWAT is closely associated with growing hair follicles and secretory coils of sweat glands. Additional abbreviations: IRS – inner root sheath; ORS – outer root sheath; DP – dermal papilla; Mx – hair matrix; SCs – stem cells. (B) Schematic of “average” hippo skin. The epidermis is relatively thin in comparison to cetaceans and displays shallow rete ridges. The dermis lacks identifiable adipocytes. Ventral, dorsal and ear skin contains pelage hair follicles that lack associated sebaceous glands. Facial skin features vibrissae follicles that also lack sebaceous glands. Tail skin contains prominent hair follicles that have vibrissa-like morphology. Several body sites contain distinct sweat glands. (C) Schematic drawing of facial skin in common bottlenose dolphin. The epidermis is very thick and features prominent rete ridges. The dermis contains numerous adipocyte clusters. In newborn dolphin, the facial skin contains actively growing vibrissae hair follicles that lack distinct collagen capsules, ringwulst, and sebaceous glands (left). In adult dolphin, facial vibrissae degrade, forming keratin-filled pits (right). Skin has no sweat glands. (D) Skin-associated genes (excluding *KRTAP* genes) that are inactivated in some or all cetaceans and sometimes in hippos (see main text). Genes are classified based on their cell type or skin structure-specific expression and ontology as projected on the human skin diagram in A. Genes in black font are inactivated in some (*) or all cetaceans but not in hippos; genes in red font are independently inactivated in some (*) or all cetaceans and both hippos; the single gene in blue font is independently inactivated in cetaceans and pygmy hippo. Inactivated genes are based on the literature [16,24,65–67,69,70,85] and new observations reported here.

While hippos and cetaceans share reduced differentiation of the epidermis and possess an ill-defined *stratum granulosum* [10,15], the hippo epidermis also differs in a number of aspects from the cetacean epidermis. In particular, hippo epidermis is much thinner, has shallower rete ridges, and it partially preserves epidermal barrier function as hippos spend substantial time on land. These differences may explain why in comparison with cetaceans, only one EDC gene, *TCHHL1* (trichohyalin-like 1), is convergently inactivated in hippos (Figure 5). *TCHHL1* is a recently described epidermal gene that is predominantly expressed in the *stratum basale* and whose specific function in keratinocyte differentiation remains unclear [83,84]. Among suprabasal keratins, only *KRT2* and *KRT77* are independently inactivated in hippos (Figure 5), and the overall degree of suprabasal keratin inactivation in hippos is less than in obligately-aquatic manatees, which are more similar to cetaceans [16]. We also show that other important epidermal genes that were reported as inactivated in cetaceans are intact in hippos. These include terminal keratinocyte differentiation-associated caspase *CASP14* [65], *PSORS1C2* (psoriasis susceptibility 1 candidate 2) [85], desmosome proteins *DSG4* and *DSC1*, transglutaminase *TGM5*, and the atypical lipoxygenase *ALOXE3* [86]. In summary, cetaceans and hippos both have inactivated copies of EDC and suprabasal keratin genes, but many more genes are knocked out in cetaceans.

Our genome-wide screen identified two new epidermal genes, *KPRP* and *ALOX15,* that are independently inactivated in cetaceans and hippos (Figure 5). KPRP (keratinocyte proline-rich protein) is considered to be an epidermal terminal differentiation-associated protein, part of the EDC, that is normally expressed in the *stratum granulosum* [87,88] – an epidermal layer that is poorly defined in hippos [15] and absent in cetaceans [10]. A recent study has identified significant association between single-nucleotide polymorphism in human *KPRP* and atopic dermatitis, a skin condition diagnosed by the disrupted epidermal barrier function [89]. Studies in *Kprp^+/−^* mutant mice further showed that KPRP haploinsufficiency leads to defective epidermal desmosome structure, a higher than normal detachment rate of differentiated keratinocytes from the *stratum corneum,* increased transepidermal water loss, and higher susceptibility to skin inflammation in experimental assays [89]. ALOX15 (arachidonate 15-lipoxygenase) is a member of the lipoxygenase family of enzymes that catalyze synthesis of bioactive lipids, including resolvins [90,91]. As the name suggests, resolvins regulate resolution of excessive inflammatory responses and can act on diverse cell types. ALOX15 is highly expressed in mouse epidermis and *Axol15*^−/−^ mutant mouse studies suggest that it is important for proper epidermal barrier function. Mutant mice develop a thickened epidermis, expanded expression of *stratum basale* markers, and increased transepidermal water loss [92]. Both genes are again independently lost in cetaceans and hippos.

We next focused our analysis on genes normally associated with skin appendages, hair follicles, sebaceous glands, and sweat glands. We confirm previous findings that some or all cetaceans have inactivated keratins of the hair inner root sheath: *KRT25, KRT26, KRT27, KRT28, KRT71, KRT72, KRT73* and *KRT74*; keratins of hair, nail and tongue papillae: *KRT32, KRT33A, KRT33B, KRT34, KRT35, KRT38, KRT39, KRT40, KRT82, KRT83* and *KRT84,* as well as keratins *KRT3* and *KRT6B* [16,93]. Of these, hippos have independently inactivated only *KRT26.* Preservations of most hair and nail keratins in hippos is consistent with the fact that unlike cetaceans, hippos have prominent keratinized hoofs [94] and distinct pelage and vibrissa hairs, albeit only in restricted anatomical sites. Particularly, brush-like hairs on hippo tail aid in spreading feces upon defecation, a behavior used by both river hippo and pygmy hippo for territorial marking [95]. Our analyses validate that cetaceans have inactivations in numerous genes associated with sebaceous gland function [69, 70]. Cetaceans are characterized by inactivations in *AWAT1, DGAT2L6, FABP9, ELOVL3, MOGAT3* and *MC5R.* We also show that *AWAT1* is inactivated in both hippos; yet unlike Lopes-Marques et al. [70], we show that these species retain an intact copy of *MOGAT3* (Figure 4). Finally, we found that the ATP-binding cassette transporter *ABCC11*, which is associated with axillary sweat gland function in humans, is inactivated in all cetaceans and independently in pygmy, but not river hippo. Both river and pygmy hippos have active sweat glands that produce pigmented secretion; therefore, we speculate that ABCC11 function is not critical for this unique aspect of hippos’ sweat gland biology.

### The Timing of Gene Inactivations in Hippopotamidae and Cetacea

To understand when gene losses occurred, we performed dN/dS analyses with the coding remnants of these genes and equations from Meredith et al. [96]. We focused on genes that were independently inactivated in Cetacea and Hippopotamidae plus two other selected genes (*MOGAT3* and *ABCC11*). To obtain robust inactivation dates, we calculated dates using eight different combinations of codon frequency model (CF1, CF2), fixed versus estimated ω values for the pseudogenic branch category, and one versus two rates for synonymous substitutions (Supplemental Table S3). Mean inactivation dates based on these estimates are shown in Figure 3 and Table 2. Mean estimates for eight genes (*ALOX15*, *AWAT1*, *KPRP*, *KRT2*, *KRT26*, *KRT77*, *KRTAP7-1*, *TCHHL1*) that were inactivated on the stem Hippopotamidae branch range from 53.92 to 5.42 Ma. These inactivation dates suggest that derived changes in hippopotamid skin, including the loss of sebaceous glands, have a long history that encompasses the entirety of the stem hippopotamid branch. The mean date for inactivation of these eight genes is ~30.5 Ma, which is near the midpoint of the stem hippopotamid branch. In addition, *ABCC11* is intact in *Hippopotamus amphibius* but appears to have been inactivated very recently in *Choeropsis liberiensis* (Figure 3; Table 2).

**Table 2.**
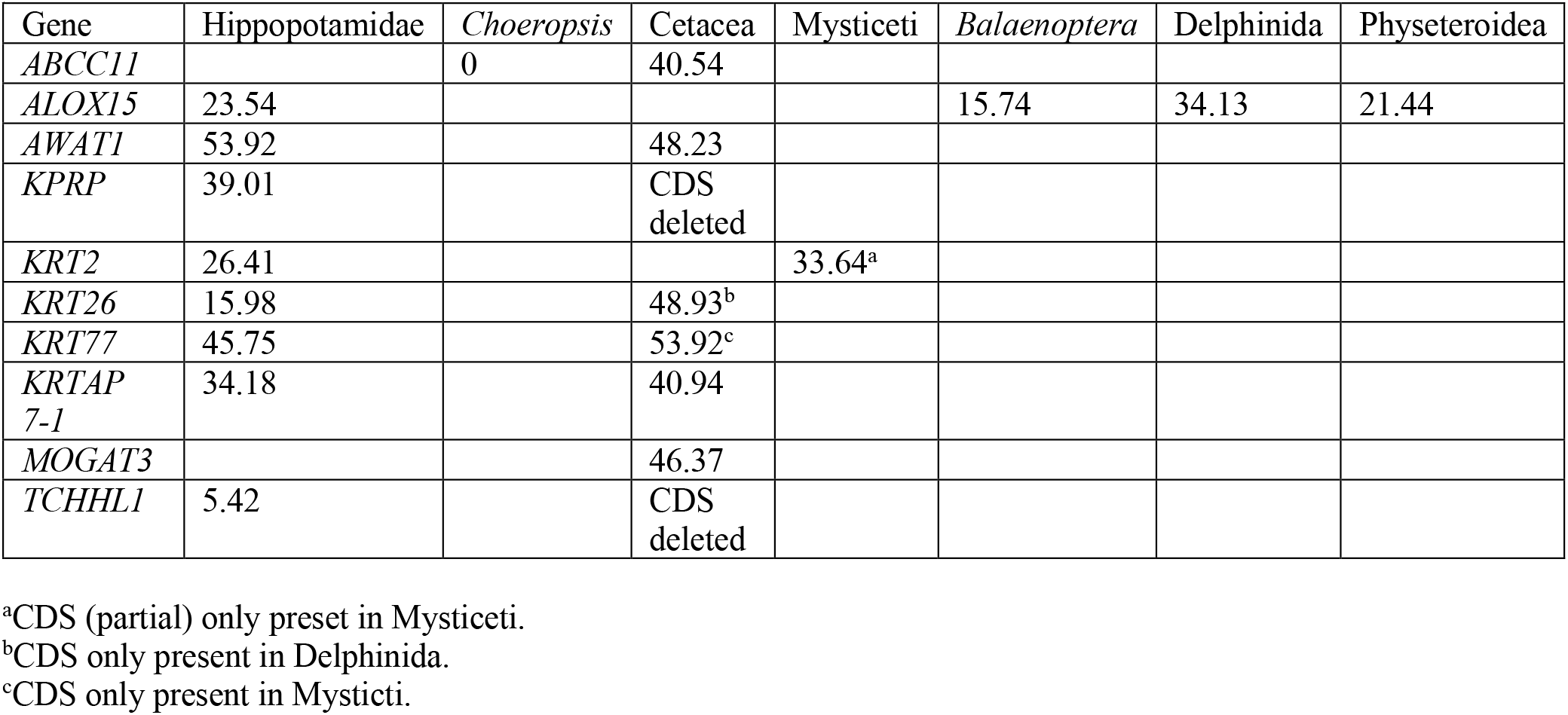
Mean inactivation dates for skin-related genes in cetaceans and hippos based on eight different combinations of codon frequency model (CF1, CF2), fixed versus estimated dN/dS values for the pseudogenic branch category, and one versus two rates for synonymous substitutions. Estimates based on individual analyses are provided in Table S3.

In the case of Cetacea, estimated inactivation dates for four genes (*ABCC11*, *AWAT1*, *KRTAP7-1*, *MOGAT3*) with clear evidence of pseudogenization on the stem cetacean branch based on shared inactivating mutations range from 48.23 to 40.54 Ma (mean = 44.02 Ma). Three additional genes are inactivated in all cetaceans, but are completely absent from either odontocetes (*KRT2, KRT77*) or mysticetes (*KRT26*). As complete gene deletion could have erased smaller mutations that inactivated these genes earlier in evolution, we used the remnants of these genes to estimate whether loss may have occurred on the stem Cetacea branch. Inactivation dates suggest that two of these genes were pseudogenized on the cetacean stem branch (53.92 Ma [*KRT77*], 48.93 Ma [*KRT26*]) whereas the date for *KRT2* [33.64 Ma] is slightly younger than the most recent common ancestor of Cetacea at ~36.72 Ma. The mean date for the six genes (*ABCC11*, *AWAT1*, *KRT26*, *KRT77*, *KRTAP7-1*, *MOGAT3*) inactivated on the stem cetacean branch is ~46.5 Ma, which is near the midpoint of the stem cetacean branch. Overall, the mean inactivation date for these six genes in cetaceans is ~16.0 million years older than the mean inactivation date for eight genes that were inactivated on the stem hippo branch. Among four overlapping genes (*AWAT1*, *KRT26*, *KRT77*, *KRTAP7-1*) with inactivation dates on the stem cetacean and stem hippo branches, the mean date for stem Cetacea (48.00 Ma) is 10.36 Ma older than the mean date for stem hippos (37.46 Ma). Two additional genes (*KPRP*, *TCHHL1*) are completely absent in all cetacean genomes that we examined, so the timing of these gene losses on the stem cetacean branch could not be estimated with our methods. Finally, *ALOX15* appears to have been independently inactivated in the common ancestor of Delphinida (34.13 Ma), on the terminal branch leading to *Physeter* (21.44 Ma), and in the common ancestor of *Balaenoptera acutorostrata* and *B. bonaerensis* (15.74 Ma) within Mysticeti (*ALOX15* is intact in *Eschrichtius robustus* and *Balaena mysticetus*).

### Integration of Molecular, Histological, and Paleontological Data

Our integrative analysis sheds new light on the key question pertaining to the evolution of Cetancodonta: Did shared morphological and behavioral features associated with aquatic and semiaquatic lifestyles evolve in the common ancestor of this clade or did they evolve independently in Cetacea and Hippopotamidae (Figure 1)? If aquatic features of hippos represent an intermediate condition in the transition from land to sea in the ancestry of Cetacea [1], behavioral study of extant hippos might provide direct insights into the behavior of the earliest cetaceans form the Eocene. Shared morphological features include the general reduction in density or complete loss of pelage hairs that cover the body, and the loss of sebaceous glands. Previous histological studies that reported the absence of sebaceous glands were inconclusive as they were limited to skin from the trunk, neck, and limbs of *Hippopotamus amphibius* that have few to no hairs [15]. Sebaceous glands normally form as part of embryonic hair follicle development. Together, sebaceous glands and hair follicles constitute an anatomically connected pilosebaceous unit. Therefore, definitive determination of the presence or absence of sebaceous glands requires a thorough investigation of hair-bearing skin in different regions of the body. We examined several regions of haired skin and provide more definitive evidence for the absence of sebaceous glands around hair follicles than previous studies [15]. We also extended these findings to a second hippopotamid species, *Choeropsis liberiensis*. Meibomian glands, which are modified sebaceous glands in the eyelid skin, are also absent in *H. amphibius*, as has also been reported for *C. liberiensis* [39]. Further, examined vibrissa follicles in newborn *Tursiops truncatus* and adult *Eschrichtius robustus* also lack sebaceous glands. These findings make a strong case for the complete body-wide absence of sebaceous glands in hippopotamids and cetaceans. The most parsimonious interpretation for the loss of sebaceous glands in hippos and cetaceans is that these glands were lost in the common ancestor of Cetancodonta. Similarly, the most parsimonious interpretation of body hair follicle reduction (hippos) or loss (cetaceans) is that pelage density started to decrease in the common ancestor of hippos and cetaceans. At the same time, the distribution of lipids is intracellular in the highly modified *stratum corneum* of cetaceans whereas hippos have intercellular *stratum corneum* lipids, which suggests parallel evolution of the highly modified epidermis in these taxa.

If pelage reduction and the loss of sebaceous glands occurred in the common ancestor of Cetancodonta, then we should expect to find evidence of shared inactivating mutations in one or more skin-specific genes in hippos and cetaceans. By contrast, the independent origins hypothesis predicts that we should see convergent gene inactivations in cetaceans and hippos. Our genomic screens identified several skin-specific genes that are inactivated in hippos and cetaceans. Strikingly, none of these genes have inactivating mutations that are shared by hippos and cetaceans. The absence of shared inactivating mutations at multiple gene loci suggests that pelage reduction and the loss of sebaceous glands occurred independently in Hippopotamidae and Cetacea.

Mean dates for the inactivation of skin genes in these two clades serve as a proxy for phenotypic changes. These dates suggest that pelage reduction/loss, the loss of sebaceous glands, and changes to the keratinization program occurred ~16.0 million years earlier in Cetacea (46.5 Ma) than in Hippopotamidae (30.5 Ma) based on all estimates for gene inactivations or ~10.5 million years earlier in Cetacea (48.0 Ma) than in Hippopotamidae (37.5 Ma) based on an overlapping set of four genes (Figure 3, Table 2). The mean date of ~48.0-46.5 Ma for Cetacea is older than the first obligately aquatic cetaceans in the family Basilosauridae (e.g., *Basilosaurus* at ~41 Ma) and instead corresponds with the oldest protocetids (e.g., *Rodhocetus*) from the Lutetian (47.8-41.3 Ma), which may have utilized both land and water as sea lions do today [97]. The mean inactivation date of ~37.5-30.5 Ma for hippopotamid skin genes, in turn, is the range of early bothriodontine anthracotheres (37.2-33.9 Ma) that are inferred to be the oldest semiaquatic members of the family Anthracotheriidae [36]. It is also noteworthy that far more skin-related genes have been inactivated in Cetacea than in Hippopotamidae (Figure 5), which is consistent with the more complete reorganization of the epidermis and its derivatives in cetaceans relative to hippopotamids (Table 1; Figures 2 and 5). Finally, our estimates of individual gene inactivations span tens of millions of years for both stem + crown cetaceans and stem + crown hippos (Figure 3, Table 2), suggesting that macroevolutionary changes to aquatic and semiaquatic habitats in these clades, respectively, have long, stepwise histories. Indeed, *AWAT1* inactivation on the stem hippopotamid branch and *ABCC11* inactivation on the *Choeropsis liberiensis* branch have estimated pseudogenization dates that are separated by ~54 million years.

Paleontological evidence also provides an opportunity to evaluate competing hypotheses that aquatic/semiaquatic features of cetaceans and hippos were acquired independently rather than in the common ancestor of this clade (Figure 1). If the initial shift to a semiaquatic lifestyle occurred in the common ancestor of Cetancodonta, then we might expect to find morphological and/or geochemical evidence for this transition in (1) stem cetancodontans that are close to the crown group, (2) the earliest branching stem cetaceans, and (3) the earliest stem hippopotamids. In the case of stem cetaceans, the most primitive and earliest branching clade is Raoellidae, which includes *Indohyus*. Thewissen et al. [98] reported that *Indohyus* has a thickened medial wall in the auditory bulla known as the involucrum. They inferred that the involucrum is associated with underwater hearing and that its presence in *Indohyus* provides evidence that the most primitive stem cetaceans were already semiaquatic. Dense limb bones (for ballast) and oxygen isotopic signatures of its teeth also suggest that *Indohyus* was semiaquatic [98]. Evidence for aquatic/semiaquatic specializations in stem cetancodontans and stem hippopotamids is less forthcoming. In the case of Cetancodonta, definitive stem members of this clade have been difficult to identify owing to the uncertainty of cladistic analyses. Possible stem cetancodontans have included a variety of anthracotheres (e.g., *Anthracotherium, Siamotherium* [56]; *Elomeryx, Heptacodon, Microbunodon* [1]), an entelodontid (*Brachyhyops* [56], a cebochoerid (*Cebochoerus* [1,99]), and the enigmatic *Andrewsarchus* [56]. We are not aware of any compelling evidence for aquatic adaptations in any of these taxa. Also, there is an emerging consensus from alternative phylogenetic hypotheses that anthracotheres comprise the paraphyletic stem group that gave rise to Hippopotamidae [36,63,99]. Many anthracotheres (e.g., *Anthracotherium, Siamotherium*) appear to have been terrestrial based on oxygen isotope values whereas taxa in the subfamily Bothriodontinae have values that are consistent with a semiaquatic lifestyle [58,59,61,62]. However, the phylogenetic placement of presumed terrestrial anthracotheres as basal, stem Hippopotamidae suggests that specializations for an aquatic/semiaquatic lifestyle evolved independently in hippopotamids and cetaceans [36,63]. Thus, paleontological evidence is largely aligned with inactivating mutations in skin-related genes that favor the independent origins hypothesis.

When sister taxa share the same anatomical, physiological, or behavioral features, as is the case for aquatic features in Hippopotamidae and Cetacea, the simplest hypothesis is that these features evolved in their common ancestor. However, our results suggest otherwise and instead support the independent evolution of features that are related to the skin in hippos and cetaceans. Given that most synapomorphies for Cetancodonta are aquatic traits (e.g., hairless or nearly hairless body, lack of sebaceous glands, lack of scrotal testes, underwater parturition/nursing) [1], morphological and behavioral support is very limited for this group if these traits evolved in parallel. Along the same lines, extant pinnipeds have lost their sweet and umami taste receptors. However, loss of these receptors occurred independently in the ancestors of the reciprocally monophyletic Phocidae (seals) and Otaroidea (Otariidae [sea lions, fur seals] + Odobenidae [walruses]) rather than in the common ancestor of Pinnipedia [100]. Numerous morphological features related to raptorial feeding and hydrodynamic locomotion also appear to have evolved independently in the three extant families of Pinnipedia [101]. In Cetacea, multiple cranial and postcranial specializations for an aquatic lifestyle (e.g., shortened humerus, loss of radial tuberosity, reduction/loss of hindlimbs, posterior migration of the blowhole, cranial ‘telescoping’) evolved convergently in odontocetes and mysticetes [1,102,103]. Morphological characters preserved in fossils and pseudogenic remnants of formerly functional genes provide complementary sources of evidence for elucidating such cases of convergent or parallel evolution.

Finally, gene inactivation dates have important implications for understanding the physiology, behavior, and appearance of extinct organisms [1,104]. In the case of cetaceans, inactivation dates for *AWAT1* and *MOGAT3* are both older than the inactivation date for *ABCC11*, which suggests that sebaceous glands were lost before sweat glands on the stem cetacean branch. Given the timing of gene inactivations for these three genes, extinct protocetid whales may have retained sweat glands but not sebaceous glands. Similarly, the hair inner root sheath keratin *KRT26* was lost relatively early on the stem cetacean branch (~49 Ma), suggesting that the program for generating body pelage hair had already been compromised at this early stage in cetacean evolution. Protocetids, which comprise a paraphyletic grade, were probably the earliest trans-oceanic cetaceans, but also spent time on land where they may have given birth and nursed their young [97,105,106]. These forms would certainly have looked different than modern whales due to their more prominent hindlimbs and primitive cranial morphology, but in other respects, such as having a largely hairless body, may have been similar to modern cetaceans.

In summary, by integrating new histological data with comprehensive analyses of molecular changes, we found strong support for the hypothesis that aquatic adaptations of the skin evolved independently in hippos and cetaceans. Our study further illustrates the potential of genomic data and in particular remnants of once functional genes as dateable ‘molecular vestiges’ to complement morphological data in providing novel insights into ancestry and timing of key trait changes and macroevolutionary transitions [68,86,96,104,107–111].

## Acknowledgments

This work was supported by NSF grant DEB-1457735 (J.G., M.S.S.), NIH grants U01-AR073159 and P30-AR075047 (M.V.P.), NSF grant DMS-1951144 (M.V.P.), and Pew Charitable Trust LEO Foundation (M.V.P.). C.F.G-J. was supported by UC Irvine Chancellor’s ADVANCE Postdoctoral Fellowship Program, NSF-Simons Postdoctoral Fellowship, NSF grant (DMS1763272, to Qing Nie), Simons Foundation Grant (594598, to Qing Nie), and by a kind gift from the Howard Hughes Medical Institute Hanna H. Gray Postdoctoral Fellowship Program. M.H. was supported by the Max Planck Society and the LOEWE-Centre for Translational Biodiversity Genomics (TBG) funded by the Hessen State Ministry of Higher Education, Research and the Arts (HMWK). We thank the Smithsonian Institution for skin samples of hippos. C. Buell provided images of mammals.

## STAR Methods

### Skin Sample Collection

The sources for skin samples used in histological analyses are as follows: *Choeropsis liberiensis* (pygmy hippopotamus) skin samples are from Smithsonian National Museum of Natural History – specimen number 395848 (unknown gender, neonatal); *Hippopotamus amphibius* (river hippopotamus) skin samples are from Smithsonian National Museum of Natural History – specimen number 254870 (male, neonatal); *Tursiops truncatus* (common bottlenose dolphin) skin samples are from Southwest Fisheries Science Center (NOAA) specimen numbers KKS0032 (unknown gender, neonatal) and KXD0206 (late term fetus); *Eschrichtius robustus* (gray whale) skin samples are from Southwest Fisheries Science Center (NOAA) Fisheries – specimen number NEB0083 (unknown gender, adult).

### Histology

Formalin-fixed *Choeropsis liberiensis* and *Hippopotamus amphibius* skin samples were first hydrated and rinsed in 1X PBS. Samples were then dehydrated through an ethanol gradient (from 25% to 100%), processed through histoclear and embedded in paraffin. Each hydration and dehydration step lasted for 12 hours. Tissues were sectioned at a thickness of 10 μm with a microtome (Leica). Samples were stained with hematoxylin and eosin using standard methods with minor modifications. Tissue sections were mounted with Permount mounting media and visualized with Nikon Ti-E Upright microscope. Tissue whole-mounts were captured with a Nikon dissecting microscope. Individual fields of *Eschrichtius robustus* rostral skin were visualized and stitched together with Keyence microscope.

### Quantification of Epidermal Thickness

Epithelial thickness of *Choeropsis liberiensis*, *Hippopotamus amphibius*, *Tursiops truncatus* and *Eschrichtius robustus* rostral skin was quantified using ImageJ (NIH). We included measurements ranging from: a) top to start of rete ridge; b) start to end of rete ridge; and c) entire epidermis including rete ridge. Up to 10 individual measurements were included per image field per species. Measurements are reported as average epidermal thickness (μm) ± standard deviation (μm) in Table 1.

### Pygmy Hippopotamus Genome

Genomic DNA from *Choeropsis liberiensis* (pygmy hippopotamus) was provided by G. Amato (formerly at New York Zoological Society). DNA was sonicated at the University of California, Riverside (UCR), Genomics Core Facility into ~550 bp fragments. We constructed a genomic library using Illumina’s NeoPrep Library Prep System. Paired-end sequencing (150 bp) was performed at UCR. Raw reads have been deposited at NCBI (accession XXXXXXXX).

### Genomic Screens for Inactivated Genes

We used a genome alignment of placental mammals with the human hg38 assembly as the reference [112] and gene loss data generated by our gene loss detection method [68,113] to screen for genes that exhibit inactivating mutations in the genomes of bottlenose dolphin, killer whale, sperm whale, common minke whale, and river hippo. We started with 19,769 genes annotated by Ensembl (http://www.ensembl.org) version 90 [114] in the human genome and considered 18,363 genes that are present in the assemblies of at least 31 of 63 placental mammals. To identify genes that were potentially inactivated on the branch leading to hippos and cetaceans, we further extracted genes that are inactivated in all cetaceans and the river hippo. We excluded genes that are intact in less than three of six terrestrial outgroup artiodactyls included in the screen (*Bos taurus* [cow], *Capra hircus* [goat], *Camelus ferus* [wild Bactrian camel], *Pantholops hodgsonii* [Tibetan antilope], *Bison bison* [bison], *Vicugna pacos* [alpaca]). Finally, we used a more recent assembly of the river hippo genome (GCA_004027065.2) to exclude instances where assembly errors mistakenly led to genes classified as inactivated in the river hippo. This resulted in a final list of 38 genes (Supplemental Table S1).

### BLAST Searches, Alignments, and Inactivating Mutations

Genomic sequences encoding ten genes of interest (*ABCC11*, *ALOX15*, *AWAT1*, *KPRP*, *KRT2*, *KRT26*, *KRT77*, *KRTAP7-1*, *MOGAT3*, *TCHHL1*) were downloaded from NCBI for *Homo sapiens* (human), *Bos taurus* (cow), and *Equus caballus* (horse). Sequences for each gene were aligned and exon annotations in *Bos* and *Equus* were compared against those in *Homo* to ensure that orthologous regions were annotated. Protein-coding sequences and flanking introns from *Bos* and *Equus* were employed in BLAST searches against other cetartiodactyls and perissodactyls, respectively, in NCBI’s ‘RefSeq Genome’ and ‘Whole-genome shotgun contigs’ databases. Additional perissodactyls included *Ceratotherium simum* (white rhinoceros) and *Dicerorhinus sumatrensis* (Sumatran rhinoceros). Additional cetartiodactyls included two camelids (*Camelus ferus* [wild Bactrian camel], *Vicugna pacos* [alpaca]), one suid (*Sus scrofa* [pig]), two bovids (*Bubalus bubalis* [water buffalo], *Capra hircus* [goat]), two giraffids (*Giraffa camelopardalis* [giraffe], *Okapia johnstoni* [okapi]), two cervids (*Axis porcinus* [hog deer], *Odocoileus virginianus* [white-tailed deer]), one hippopotamid (*Hippopotamus amphibius* [river hippopotamus]), four mysticetes (*Balaena mysticetus* [bowhead, downloaded from http://www.bowhead-whale.org/], *Balaenoptera acutorostrata* [common minke whale], *B. bonaerensis* [Antarctic minke whale], *Eschrichtius robustus* [gray whale]), and ten odontocetes (*Physeter macrocephalus* [sperm whale], *Lipotes vexillifer* [baiji], *Delphinapterus leucas* [beluga], *Phocoena phocoena* [harbor porpoise], *Neophocaena asiaorientalis* [narrow-ridged finless porpoise], *Orcinus orca* [killer whale], *Lagenorhynchus obliquidens* [Pacific white-sided dolphin], *Sousa chinensis* [Indo-Pacific humpback dolphin], *Tursiops aduncus* [Indo-Pacific bottlenose dolphin], *Tursiops truncatus* [common bottlenose dolphin]). Additional searches were performed with other perissodactyls or cetartiodactyls when the initial searches with *Equus* and *Bos* were unsuccessful in retrieving complete orthologs. Megablast was employed for highly similar sequences and blastn for less similar sequences. Complete protein-coding sequences and intervening introns were imported into Geneious 11.1.5 [115] and aligned against reference sequences with MAFFT [116] with minor adjustments by eye. Aligned sequences were annotated for exons and inspected for splice site mutations. Illumina sequences for *Choeropsis liberiensis* were imported into Geneious and protein-coding sequences for the above-mentioned genes were obtained using a map to reference approach with probe sequences from the closely related *Hippopotamus amphibius*. We allowed for a maximum mismatch of 6% per read and required at least two reads for base calling with a consensus threshold of 65%. We also used MAFFT to align complete protein-coding sequences from all taxa for each gene.

### Inactivating Mutations

Final alignments for protein-coding sequences for each gene included 15-29 taxa given that some genes are deleted in one or more cetaceans. We inspected the final protein-coding alignment for each gene for inactivating mutations including exon deletions, frameshift insertions and deletions, altered start and stop codons, and premature stop codons (splice site mutations screened above). Parsimony optimizations with delayed transformation (deltran) were performed with PAUP* 4.0a150 [117]to map inactivating mutations to branches of the species tree (see below) for each gene.

### Phylogenetic Analyses

RAxML 8.2.11 [118] was run in Geneious to estimate maximum likelihood gene trees for each protein-coding alignment. Gene trees were inspected for suspicious relationships that conflict with the species tree (e.g., *Camelus* grouping with *Sus* instead of *Vicugna*) but none were found. Instead, gene tree incongruence was confined to conflicts that are readily explained by ILS such as Ruminantia grouping with Suoidea or *Physeter* grouping with Mysticeti [119]. Rapid bootstrap analysis (100 pseudoreplications) and a search for the best tree were performed in a single run. These analyses were performed with a GTR + Γ model of sequence evolution.

### DN/dS Analyses

DN/dS analyses were performed with the codeml program of PAML 4.4 [120]. Analyses for each gene were performed with separate dN/dS categories for functional branches that lack inactivating mutations, fully pseudogenic branches that post-date the occurrence of an inactivating mutation on an earlier branch, and each transitional branch that records the first occurrence of an inactivating mutation (e.g., [96]). Analyses were performed with the codon frequency 1 (CF1) and codon frequency 2 (CF2) models of codeml. We also performed analyses with estimated and fixed (dN/dS =1.00) values for the fully pseudogenic branch category. We employed a species tree with higher level (interordinal, interfamilial) relationships from Meredith et al. [121] and intrafamilial relationships from Hassanin et al. [122] for terrestrial cetartiodactyls and McGowen et al. [123] for cetancodontans.

### Estimation of Gene Inactivation Times

Equations from Meredith et al. [96] were used to estimate gene inactivation times in hippopotamids and cetaceans. We performed calculations using eight different combinations of codon model (CF1 or CF2), fixed (1.0) versus estimated values for dN/dS on fully pseudogenic branches, and equations that allow for one versus two synonymous substitution rates [96]. Mean inactivation dates for each gene are averages based on these eight different combinations. Divergence times for relevant nodes in these calculations were taken from McGowen et al. [123].

## Supplemental Tables

**Table S1.** List of genes that are inactivated in cetaceans and hippopotamids.

**Table S2.** Inactivating mutations in cetaceans and hipppotamids.

**Table S3.** Estimated inactivation dates for pseudogenized skin genes.

